# Less physical activity and more varied and disrupted sleep is associated with a less favorable metabolic profile in adolescents

**DOI:** 10.1101/2020.01.31.928333

**Authors:** Vaka Rognvaldsdottir, Robert J. Brychta, Soffia M. Hrafnkelsdottir, Kong Y. Chen, Sigurbjorn A. Arngrimsson, Erlingur Johannsson, Sigridur L. Guðmundsdottir

**Affiliations:** Center of Sport and Health Sciences, University of Iceland, Reykjavik, Iceland; Diabetes, Endocrinology and Obesity Branch, National Institute of Diabetes and Digestive and Kidney Diseases, Bethesda, MD, USA; Department of Sport and Physical Activity, Western Norway University of Applied Sciences, Bergen, Norway

## Abstract

**Background:** Sleep and physical activity are modifiable behaviors that play an important role in preventing overweight, obesity, and metabolic health problems. Studies of the association between concurrent objective measures of sleep, physical activity, and metabolic risk factors among adolescents are limited.

**Objective:** The aim of the study was to examine the association between metabolic risk factors and objectively measured school day physical activity and sleep duration, quality, onset, and variability in adolescents.

**Materials and Methods:** We measured one school week of free-living sleep and physical activity with wrist actigraphy in 252 adolescents (146 girls), aged 15.8±0.3 years. Metabolic risk factors included body mass index, waist circumference, total body and trunk fat percentage, resting blood pressure, and fasting glucose and insulin levels. Multiple linear regression adjusted for sex, parental education, and day length was used to assess associations between metabolic risk factors and sleep and activity parameters.

**Results:** On average, participants went to bed at 00:22±0.88 hours and slept 6.2±0.7 hours/night, with 0.83±0.36 hours of awakenings/night. However, night-to-night variability in sleep duration (0.87±0.57 hours) and bedtime (0.79±0.58 hours) was considerable. Neither average sleep duration nor mean bedtime was associated with any metabolic risk factors. However, greater night-to-night variability in sleep duration was associated with higher total body (β=1.9±0.9 %/h, p=0.03) and trunk fat percentage (β=1.6±0.7 %/h, p=0.02), poorer sleep quality (more hours of awakening) was associated with higher systolic blood pressure (β=4.9±2.2 mmHg/h, p=0.03), and less physical activity was associated with higher trunk fat percentage (p=0.04) and insulin levels (p=0.01).

**Conclusion:** Greater nightly variation in sleep, lower sleep quality, and less physical activity was associated with a less favorable metabolic profile in adolescents. These findings support the idea that, along with an adequate amount of sleep and physical activity, a regular sleep schedule is important to the metabolic health of adolescents.

## Introduction

The prevalence of overweight in the world has nearly tripled from 1975-2016, with over 39% of adults and 18% of children and adolescents being overweight or obese [1]. Greater total body and central adiposity is associated with increased risk of cardio-metabolic comorbidities, such as hypertension and diabetes [2, 3]. Prevalence of metabolic syndrome is high among obese children and adolescents and increases with higher central obesity [4]. Along with diet, sleep and physical activity have been identified as important modifiable risk factors implicated in the development of overweight, obesity, and metabolic health problems [5].

The importance of adequate sleep for health and daily functioning in adolescents is well established [6, 7], although most studies are based on subjective data. Most national and international guidelines focus on recommendations for sleep duration, since prior research has demonstrated that insufficient sleep duration during adolescence is associated with a variety of cognitive, psychological, and health risks, including higher body mass index (BMI) [8–10], greater body fat [11], and increased insulin resistance [12]. However, emerging evidence suggests that sleep quality [13–15] and timing may also affect adolescent cardiometabolic risk factors [7]. For instance, later bedtimes are associated with greater BMI [10, 16], body fat [11] and higher systolic blood pressure in children and adolescents [17]. Markers of irregular sleep schedules, such as high variability in sleep duration or greater shifts in sleep timing and duration on weekends, have also been associated with greater adiposity and abdominal obesity [18, 19] and higher BMI and insulin levels [20] in children and adolescents. Studies also suggest that long-term exposure to a disrupted sleep schedule [21] or low physical activity [22] can increase the risk of developing metabolic syndrome, while higher levels of physical activity in children and adolescents are associated with favorable body mass index, lower adiposity, and better cardio-metabolic health [23, 24]. However, since studies with simultaneous objective measures of sleep, activity, and metabolic risk factors are sparse, it is not known whether physical activity and sleep contribute to a better cardiometabolic profile independently.

The aim of the study was to examine associations between metabolic risk factors and concurrent objective measures of free-living sleep and physical activity among Icelandic adolescents. We hypothesized that less activity, shorter sleep duration, poorer sleep quality, and more varied sleep schedules will associate with less favorable cardiometabolic profiles.

## Methods

### Study design and data collection

All students attending the second grade in six of the largest primary schools in Reykjavik, Iceland were invited to participate in a longitudinal cohort studying health, cardiovascular fitness, and physical activity initiated at seven to eight years of age (N = 320, 82% participated) [25]. At the age of 15, concurrent measurement of sleep and activity with wrist activity was introduced [26] and students from the cohort and all others enrolled in the same grade at the respective schools were invited to participate (N = 411); 315 agreed (response rate 77%), and 252 had complete data for questionnaire, body composition, sleep, and physical activity measurements. Two participants were missing waist circumference and blood pressure measurements and 13 refused blood draws for serum glucose and insulin. Study participation is shown in Figure 1.

**Figure 1.**
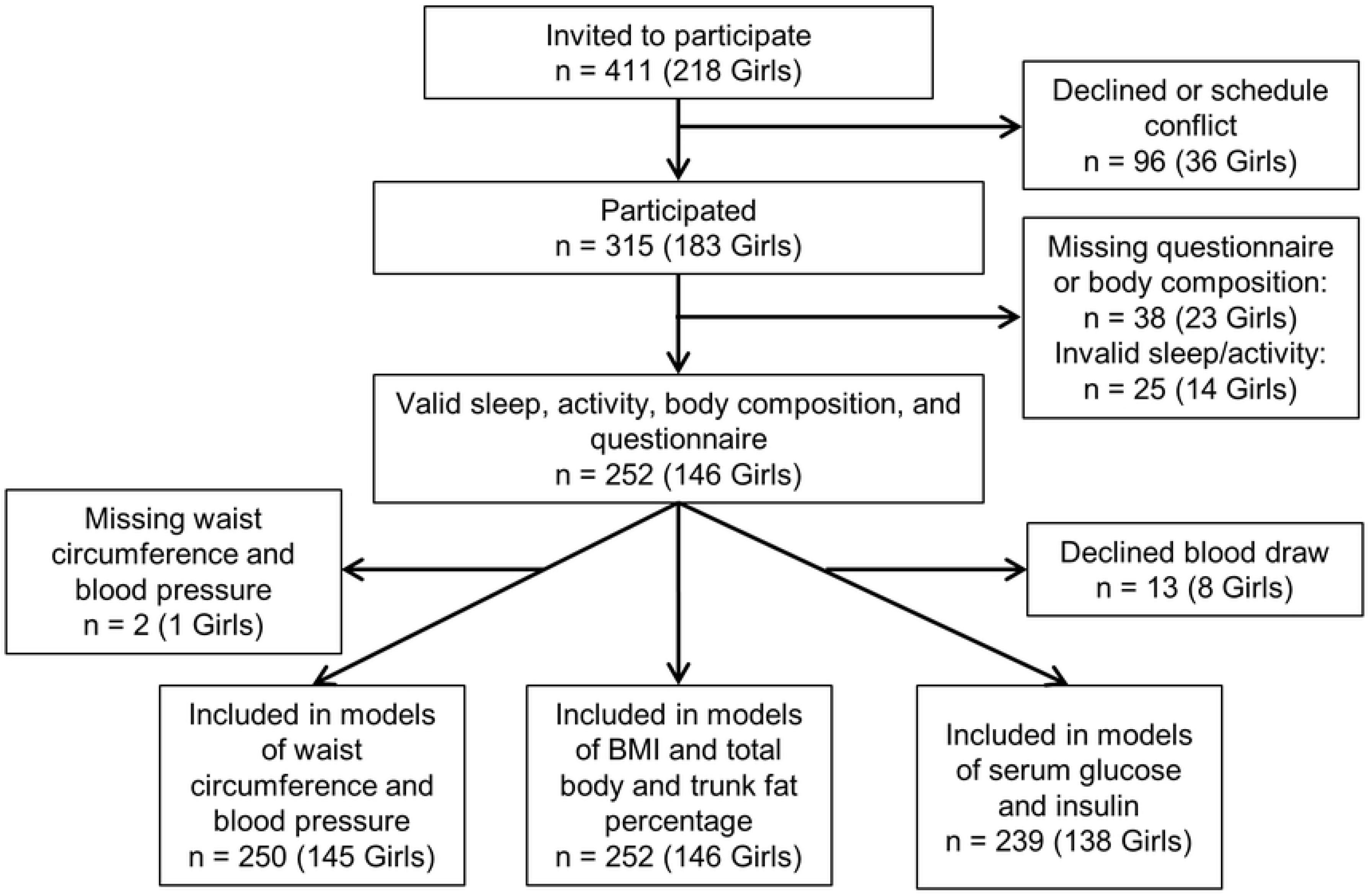
Flow chart describing study participation.

Written informed consent was obtained from all participants and their guardians. The study was approved by the National Bioethics Committee, the Icelandic Data Protection Authority (Study number: VSNb2015020013/13.07), and the Icelandic Radiation Safety Authority. The study was conducted in agreement with the guidance provided in the Declaration of Helsinki.

### Sleep and physical activity parameters

Free-living sleep and physical activity were measured with a wrist-worn accelerometer (GT3X+, Actigraph Inc., Pensacola, FL, USA). The accelerometer was placed on the non-dominant wrist at school and the participant was asked to wear it continuously for a week. Physical activity counts and sleep duration, timing, and quality, were computed with Actilife software version 6.13.0 (Actigraph). The Sadeh algorithm, validated for adolescents [27], was used to detect sleep onsets and awakenings, which were visually inspected and adjusted as necessary by two expert scorers based on daily sleep logs maintained during the week of actigraphy. Wear time and vector magnitude of physical activity counts were computed in MATLAB (version R2013a MathWorks, Natick, MA, USA) using previously described algorithms [28, 29]. Since only one week of accelerometer data was collected, we focused our analysis on school days (Monday-Friday) and nights (Sunday-Thursday) and did not include data for weekends or holidays. Participants with ≥3 school days and wear time ≥14 hours, were included in the analyses. Sleep and activity parameters are defined in Table 1.

**Table 1.** Sleep and physical activity parameters.

| Actigraphy parameter | Definition |
| --- | --- |
| Sleep duration | Time spent asleep between sleep onset and awakening (hours/night) |
| Variability in sleep duration | Night-to-night variation (standard deviation) of sleep duration (hours) |
| Bedtime | Time of sleep onset (clock time) |
| Variability in bedtime | Night-to-night variation (standard deviation) of sleep onset (hours) |
| WASO | Wake time after sleep onset (hours/night) |
| Physical activity | Activity/wear time (3D-counts/minutes of wear/day) |

### Body composition

Height, weight, and waist circumference was measured at participants’ schools. Standing height was measured with a stadiometer (Seca model 217, Seca Ltd. Birmingham, UK) to the nearest 0.1 cm. Body weight was measured to the nearest 0.1 kg using a scale (Seca model 813, Seca Ltd. Birmingham, UK) with participants wearing light clothes. BMI was calculated by dividing weight by height squared (kg/m^2^). Waist circumference was measured to the nearest 0.1 cm using a tape measure next to skin at the narrowest place between the lowest rib and the iliac crest. Fat-free and fat mass were measured with dual-energy X-ray absorptiometry (DXA) using a GE LUNAR scanner (General Electric Lunar iDXA) at the Icelandic Heart Association. All DXA measurements were performed by a certified radiologist. Body fat percentage was calculated by dividing total fat mass by the total body mass (fat mass + lean mass + bone mineral content) and trunk fat percentage was calculated by dividing the total trunk fat mass by total trunk mass. Resting blood pressure was measured on the left arm of seated participants and the average of three measurements was used for analysis.

### Serum measures

Fasting blood samples were obtained using standard procedures after overnight fasting; samples were analyzed for glucose and insulin. Insulin (mU/L) in serum was measured using the INSULIN assay from Roche, a sandwich electrochemiluminescence immunoassay ECLIA on Cobas e 411 (Roche, Switzerland). The inter-assay coefficient of variation was < 5.06% using a frozen serum pool and < 2.36% using quality control samples from Roche. Glucose (mmol/L) in serum was measured using the GLUC2 assay from Roche, an enzymatic reference method with hexokinase. The measurements were done on a Cobas e 311 (Roche, Switzerland). The inter-assay coefficient of variation was <1.65% using a frozen serum pool and <1.66% using quality control samples from Roche.

### Survey questions and environmental data

Students provided the educational attainment of both mother and father from the following options (presented in Icelandic): 1 = “elementary degree”, 2 = “secondary degree”, 3 = “trade school degree”, 4 = “university degree”, 5 = “other”, 6 = “do not know”, 7 = “do not want to answer”. For the current analysis, responses were recoded into a binary variable: 1 = “parent with a university degree” or 0 = “no parent with a university degree”, as described previously [30]. Information on day length (hours of day light) was obtained from National Oceanic and Atmospheric Administration (NOAA) Earth System Research Laboratory Solar Calculator [31].

### Statistical analyses

Chi-squared tests and unpaired T-tests showed no differences in the gender distribution, parental education, age, body composition, or cardiometabolic risk markers of participants with and without complete actigraphy data (Table S1). T-test for independent samples was used to assess whether participant characteristics, sleep parameters, and physical activity differed between the sexes or by parental educational attainment. Separate multiple linear regression models adjusted for sex, parental education, and day length, were used to explore the associations of each sleep and activity parameter with body composition parameters (BMI, total body fat percentage and trunk fat percentage) and metabolic risk factors (insulin, glucose, blood pressure). In further analysis, body composition and metabolic risk factors were included as response variables while sleep duration, WASO, sleep variability, and physical activity were all simultaneously included as predictor variables in models additionally adjusted for sex, parental education, and day length. In a separate analysis, body composition and metabolic risk factors were again included as response variables while bedtime and bedtime variability were simultaneously included as predictor variables in models additionally adjusted for physical activity, sex, parental education, and day length. Statistical analyses were carried out in Rstudio (Boston, MA, USA, Version 1.1.456) using R statistical software (https://www.r-project.org/, Version 3.5.1). Statistical significance level was set at p<0.05.

## Results

Participant characteristics are shown in Table 2. Although boys were taller and heavier than girls, BMI (overall mean = 21.9±3.0 kg/m^2^) did not differ between the sexes. Overall, 87% of the participants had BMI below 25 kg/m^2^, 10% had 25≤ BMI <30 kg/m^2^, and 2.5% had BMI ≥ 30 kg/m^2^. Boys had lower total body and trunk fat percentage, smaller waist circumference, and slightly higher systolic pressure, but there were no sex differences in age, parental educational attainment, or serum insulin and glucose levels. Participants with and without a parent with a university degree did not differ in characteristics, body composition, blood pressure, or serum insulin and glucose (Table S2).

**Table 2.** Participants characteristics.

|  | All (252) | Boys (106) | Girls (146) | p (Boys vs Girls) |
| --- | --- | --- | --- | --- |
| **Subjects Characteristics** |  |  |  |  |
| Age, years | 15.8 ± 0.3 | 15.8 ± 0.3 | 15.9 ± 0.3 | 0.12 |
| Height, cm | 172.0 ± 8.0 | 178.5 ± 6.0 | 167.3 ± 5.6 | **<0.001** |
| Weight, kg | 64.8 ± 10.6 | 68.9 ± 10.3 | 61.9 ± 9.7 | **<0.001** |
| Body mass index, kg/m<sup>2</sup> | 21.9 ± 3.0 | 21.6 ± 2.9 | 22.1 ± 3.1 | 0.21 |
| Body fat, % | 25.1 ± 8.6 | 18.2 ± 6.6 | 30.2 ± 5.9 | **<0.001** |
| Trunk fat, % | 23.6 ± 9.6 | 17.0 ± 7.8 | 28.3 ± 7.8 | **<0.001** |
| Waist circumference, cm* | 70.6 ± 7.1 | 73.5 ± 6.2 | 68.6 ± 7.1 | **<0.001** |
| Diastolic pressure, mmHg* | 70.8 ± 5.5 | 70.2 ± 5.6 | 71.2 ± 5.3 | 0.21 |
| Systolic pressure, mmHg* | 115.1 ± 12.8 | 118.6 ± 9.9 | 112.6 ± 14.0 | **0.03** |
| Glucose, mmol/L** | 4.9 ± 0.5 | 4.9 ± 0.4 | 4.9 ± 0.5 | 0.57 |
| Insulin, mU/L** | 9.7 ± 4.7 | 8.4 ± 3.7 | 10.6 ± 5.2 | 0.73 |
| Parent with university degree, N (%) | 193 (76.6%) | 83 (78.3%) | 110 (75.3%) | 0.69 |

Data presented as mean ± standard deviation; \*250 participants (105 Boys, 145 Girls); \*\*239 participants (101 Boys, 138 Girls); Boldface type indicates significant difference (p<0.05).

Sleep and physical activity parameters did not differ between the sexes (Table 3) or between those with or without a parent with a university degree (Table S2). On average, participants spent 7.05 ± 0.82 hours in bed on school nights, going to bed at 00:22 ± 0.88 hours and rising at 07:27 ± 0.62 hours. While in bed, participants were awake 0.83 ± 0.36 hours and asleep 6.19 ± 0.73 hours. Night-to-night variations in bedtime and sleep duration were 0.79 ± 0.58 hours and 0.87 ± 0.57 hours, respectively.

**Table 3.** Summary sleep and physical activity parameters.

|  | **All (252)** | **Boys (106)** | **Girls (146)** | **p (Boys Vs. Girls)** |
| --- | --- | --- | --- | --- |
| Time in bed, hours/night | 7.05 ± 0.82 | 6.99 ± 0.84 | 7.09 ± 0.80 | 0.34 |
| Sleep duration, hours/night | 6.19 ± 0.73 | 6.13 ± 0.78 | 6.24 ± 0.70 | 0.28 |
| Bedtime, hh:mm ± hours | 00:22 ± 0.88 | 00:28 ± 0.90 | 00:18 ± 0.87 | 0.12 |
| Rise time, hh:mm ± hours | 07:27 ± 0.62 | 07:30 ± 0.68 | 07:24 ± 0.57 | 0.25 |
| WASO, hours/night | 0.83 ± 0.36 | 0.83 ± 0.35 | 0.83 ± 0.36 | 0.98 |
| Variability in sleep duration, hours | 0.87 ± 0.57 | 0.92 ± 0.65 | 0.84 ± 0.51 | 0.31 |
| Variability in bedtime, hours | 0.79 ± 0.58 | 0.85 ± 0.68 | 0.74 ± 0.49 | 0.17 |
| Activity, 1000 counts/minutes of wear/day | 2.21 ± 0.50 | 2.24 ± 0.47 | 2.18 ± 0.52 | 0.42 |
| Wear time, hours/night | 23.76 ± 0.39 | 23.69 ± 0.50 | 23.81 ± 0.29 | **0.03** |
| Valid days, days | 4.6 ± 0.6 | 4.5 ± 0.7 | 4.7 ± 0.5 | 0.13 |
| Day length, hours/day | 17.46 ± 1.87 | 17.35 ± 1.86 | 17.54 ± 1.88 | 0.42 |

Data presented as mean ± standard deviation; WASO: wake after sleep onset; Boldface type indicates significant difference (p<0.05).

The association of metabolic risk factors to physical activity and sleep duration, quality, and variability are shown in Table 4. Average sleep duration was not associated with body composition or metabolic parameters. However, the nightly variability of sleep duration was positively associated with both total body and trunk fat percentages. WASO, an indicator of sleep quality, was not associated with body composition but positively associated with systolic blood pressure. Physical activity was negatively associated with trunk fat percentage and fasting insulin levels. Neither physical activity nor any of the sleep parameters was associated with fasting plasma glucose. All significant associations persisted when average sleep duration, WASO, nightly sleep variability, and physical activity were included in the same model (Table 4, combined model).

**Table 4.** Association of metabolic risk factors to physical activity and sleep duration, quality, and variability.

| | Sleep duration, $\beta \pm SE$ (p) | WASO, $\beta \pm SE$ (p) | Nightly variability in sleep duration, $\beta \pm SE$ (p) | Physical activity, $\beta \pm SE$ (p) |
| --- | --- | --- | --- | --- |
| Body mass index, kg/m<sup>2</sup> |  |  |  |  |
| Individual model | -0.201 ± 0.262 (0.44) | -0.812 ± 0.529 (0.13) | 0.612 ± 0.332 (0.07) | 0.567 ± 0.394 (0.15) |
| Combined model | -0.274 ± 0.26 (0.29) | -0.888 ± 0.529 (0.09) | 0.518 ± 0.328 (0.12) | 0.542 ± 0.382 (0.16) |
| Trunk fat, % |  |  |  |  |
| Individual model | -0.675 ± 0.674 (0.32) | -2.427 ± 1.361 (0.08) | **1.854 ± 0.854 (0.03)** | **-2.057 ± 1.014 (0.04)** |
| Combined model | -0.434 ± 0.677 (0.52) | -2.189 ± 1.378 (0.11) | **2.161 ± 0.847 (0.01)** | **-2.115 ± 0.989 (0.03)** |
| Total body fat, % |  |  |  |  |
| Individual model | -0.695 ± 0.537 (0.20) | -1.885 ± 1.084 (0.08) | **1.606 ± 0.680 (0.02)** | -1.478 ± 0.808 (0.07) |
| Combined model | -0.521 ± 0.539 (0.33) | -1.722 ± 1.098 (0.12) | **1.824 ± 0.673 (0.01)** | -1.514 ± 0.789 (0.06) |
| Waist circumference, cm |  |  |  |  |
| Individual model | -0.202 ± 0.594 (0.73) | -1.729 ± 1.206 (0.15) | 1.049 ± 0.765 (0.17) | -0.422 ± 0.894 (0.64) |
| Combined model | -0.169 ± 0.587 (0.77) | -1.724 ± 1.200 (0.15) | 1.134 ± 0.752 (0.13) | -0.473 ± 0.867 (0.59) |
| Diastolic pressure, mmHg |  |  |  |  |
| Individual model | 0.202 ± 0.484 (0.68) | 0.774 ± 0.982 (0.43) | 0.156 ± 0.623 (0.80) | -0.964 ± 0.728 (0.19) |
| Combined model | 0.319 ± 0.476 (0.50) | 0.879 ± 0.978 (0.37) | 0.284 ± 0.613 (0.64) | -1.089 ± 0.701 (0.12) |
| Systolic pressure, mmHg |  |  |  |  |
| Individual model | 1.045 ± 1.095 (0.34) | **4.865 ± 2.221 (0.03)** | -1.824 ± 1.408 (0.20) | 0.835 ± 1.646 (0.61) |
| Combined model | 0.987 ± 1.086 (0.36) | **4.865 ± 2.213 (0.03)** | -2.015 ± 1.395 (0.15) | 0.642 ± 1.608 (0.69) |
| Glucose, mmol/L |  |  |  |  |
| Individual model | -0.077 ± 0.042 (0.07) | -0.022 ± 0.086 (0.80) | -0.008 ± 0.054 (0.88) | -0.026 ± 0.064 (0.68) |
| Combined model | -0.074 ± 0.041 (0.08) | -0.020 ± 0.085 (0.82) | -0.008 ± 0.053 (0.89) | -0.002 ± 0.061 (0.97) |
| Insulin, mU/L |  |  |  |  |
| Individual model | 0.015 ± 0.409 (0.97) | -0.726 ± 0.826 (0.38) | 0.558 ± 0.516 (0.28) | **-1.813 ± 0.613 (0.003)** |
| Combined model | 0.255 ± 0.410 (0.53) | -0.486 ± 0.839 (0.56) | 0.851 ± 0.513 (0.10) | **-1.901 ± 0.588 (0.001)** |

Sleep duration is in units of hours/nights; WASO: wake after sleep onset, in hours/night; Variability in sleep duration is in units of hours; Physical activity is in units of 1000 counts/minutes of wear/day; Individual models adjusted for sex, parental education, and day length; Combined models include sleep duration, WASO, nightly variability in sleep duration, physical activity, sex, parental education, and day length; Boldface type indicates significant relationships ( $p < 0.05$ ).

The association of metabolic risk factors to average bedtime and nightly bedtime variability is shown in Table 5. Mean bedtime was not associated with any of the body composition or metabolic parameters after adjusting for sex, parental education, and day length. However, using the same covariates, bedtime variability was positively associated with waist circumference and total body and trunk fat percentage and negatively associated with systolic blood pressure. All significant relationships persisted when average bedtime and nightly bedtime variability were included in a combined model, adjusted for physical activity, sex, parental education, and day length.

**Table 5.** Association of metabolic risk factors to average bedtime and nightly variability in bedtime.

|  | **Bedtime, $\beta \pm SE$ (p)** | **Nightly variability in bedtime, $\beta \pm SE$ (p)** |
| --- | --- | --- |
| Body mass index, kg/m<sup>2</sup> |  |  |
| Individual model | 0.222 ± 0.221 (0.31) | 0.622 ± 0.327 (0.06) |
| Combined model | 0.182 ± 0.220 (0.41) | 0.632 ± 0.328 (0.06) |
| Trunk fat, % |  |  |
| Individual model | 0.454 ± 0.576 (0.43) | **2.286 ± 0.844 (0.007)** |
| Combined model | 0.396 ± 0.567 (0.49) | **2.151 ± 0.843 (0.011)** |
| Total body fat, % |  |  |
| Individual model | 0.479 ± 0.458 (0.30) | **2.096 ± 0.669 (0.002)** |
| Combined model | 0.417 ± 0.450 (0.36) | **1.987 ± 0.669 (0.003)** |
| Waist circumference, cm |  |  |
| Individual model | 0.069 ± 0.501 (0.89) | **1.483 ± 0.734 (0.044)** |
| Combined model | 0.011 ± 0.501 (0.98) | **1.463 ± 0.740 (0.049)** |
| Diastolic pressure, mmHg |  |  |
| Individual model | -0.093 ± 0.407 (0.82) | -0.299 ± 0.601 (0.62) |
| Combined model | -0.046 ± 0.408 (0.91) | -0.349 ± 0.603 (0.56) |
| Systolic pressure, mmHg |  |  |
| Individual model | -1.342 ± 0.926 (0.15) | **-3.017 ± 1.359 (0.03)** |
| Combined model | -1.223 ± 0.924 (0.19) | **-2.865 ± 1.365 (0.04)** |
| Glucose, mmol/L |  |  |
| Individual model | 0.041 ± 0.035 (0.24) | 0.028 ± 0.053 (0.60) |
| Combined model | 0.041 ± 0.036 (0.26) | 0.025 ± 0.053 (0.64) |
| Insulin, mU/L |  |  |
| Individual model | 0.068 ± 0.349 (0.85) | 0.179 ± 0.523 (0.73) |
| Combined model | 0.095 ± 0.343 (0.78) | 0.062 ± 0.516 (0.91) |

Bedtime is in units of hours from midnight; Variability in bedtime is in units of hours; Individual models adjusted for sex, parental education, and day length; Combined models include bedtime, nightly variability in bedtime, physical activity, sex, parental education, and day length; Boldface type indicates significant relationships ( $p < 0.05$ ).

## Discussion

We studied the free-living sleep and physical activity patterns on school days in a sample of 15-year-old Icelandic boys and girls, and, as hypothesized, we found that greater nightly variation in sleep, lower sleep quality, and less physical activity was associated with less favorable indicators of metabolic health. Surprisingly, neither mean bedtime nor average sleep duration on school nights was associated with any of the cardiometabolic risk factors measured in our study. These findings support the idea that, along with an adequate amount of sleep and physical activity, a regular sleep schedule is important to the cardiometabolic health of adolescents.

We found that the participants in our study had considerable nightly variation in sleep duration (0.87 hours) and bedtime (0.79 hours) on school nights and that higher nightly variability in these parameters was related to greater measures of adiposity. These findings are consistent with previous studies of sleep variability and metabolic health, although to date study of this relationship in adolescents has been sparse. He et al., also found that high variability in sleep duration was associated with greater central adiposity in a group of similarly aged adolescents, even after controlling for food intake [18]. However, there were several notable differences in our regression analyses: (1) we included a measure of physical activity, a well-documented contributor to body composition and metabolic health, and (2) we excluded non-school nights of sleep, when sleep patterns are typically less regular and very different from school night for adolescents [26]. The exclusion of non-school nights from our analysis may partly explain the lower night-to-night variation in sleep duration for our participants compared to He, et. al. (0.87 hours vs. 1.2 hours) [18]. Despite these differences, we observed a similar robust association between sleep variability and adiposity. For instance, our combined regression model indicates that a 30 min increase in variability in nightly sleep duration would lead to a 1.1% increase in trunk fat and a 0.9% increase in total body fat. All else being equal, a participant in the ninetieth percentile of night-to-night sleep variability in our cohort (1.49 h over the school week) could be expected to have 2.12 percentage points higher body fat than a participant in the tenth percentile (0.33 h over the school week). The similarity between relationships with variability in sleep duration and bedtime was also not surprising, since the two measures were highly correlated (r=0.72, p<0.0001). These findings reinforce the idea that adolescents should maintain a regular sleep schedule.

Contrary to our hypothesis, we did not observe a relationship between mean bedtime or average sleep duration and metabolic risk factors. While a number of previous studies have noted a positive association between self-reported short sleep and obesity in adolescents [32], some more recent studies find a lack of this relationship while measuring sleep with actigraphy [17, 18]. In agreement with our findings, He et al found that high variability in sleep duration, but not mean sleep duration, was associated with greater central adiposity [18]. However, we did not find an association between average bedtime and blood pressure, as demonstrated by Mi, et al in a group of mostly younger adolescents (12.4 ± 2.6 y) [17]. It is also worth noting that our Icelandic cohort was generally lean, with a 12% prevalence of overweight and obesity. Thus, our results may reflect a more subtle relationship between sleep parameters and body composition than found in studies with greater prevalence of overweight and obesity.

We also found that more wakefulness during the sleep period (WASO) was associated with higher systolic blood pressure. This agrees with a previous cross-sectional analysis of healthy adolescents which reported that lower actigraphy-measured sleep quality was associated with pre-hypertension and hypertension [33]. Surprisingly, increased variability in bedtime was associated with lower systolic blood pressure. The meaning of those contradicting assciations with systolic blood pressure is unclear and needs further investigation. Pediatric systolic blood pressure is known to vary due to a variety of reasons [34] and further investigation of the effect of bedtime may elucidate the relationship between the two variables.

The positive influence of physical activity and the metabolic health of adolescents is well documented [35]. Our finding that wrist-actigraphy measured physical activity was inversely associated with trunk fat percentage and serum insulin levels is largely confirmatory of previous work and consistent with our hypothesis. To put these findings in perspective, all else being equal, one would expect those with a physical activity level 30% above the group mean (approximately the ninetieth percentile for this cohort) to have 2.3 mU/L lower fasting insulin and 2.6% lower trunk fat than those with an activity level 30% below the group mean (approximately the tenth percentile). Although these are substantial cross-sectional differences, they are smaller than the changes in these measures observed during aerobic exercise interventions in adolescents [36].

The potential causal pathways between irregular sleep patterns and increased body fat are not yet clear. Study of healthy non-overweight children (5-12 years) found that shorter sleep duration and poor sleep continuity were associated with overeating and other behavior related to obesity risk [37]. Sleep timing, duration, and quality are known to affect regulatory hormones, such as cortisol and growth hormone, as well as appetite regulatory hormones leptin and ghrelin [38]. Thus, high variability in sleep schedule may affect appetite control and contribute to greater adiposity and markers of poorer metabolic health. However, we did not have a measure of food intake and, thus, cannot explore potential relationships between diet, sleep variability, and metabolic health and, based on our cross-sectional study design, we cannot rule out reverse causality.

A strength of this study is the objective measurement of sleep patterns, physical activity, and body composition. Most previous studies of sleep in this age group have relied on self- or parent-report of typical time in bed or bed- and rise-times. Self-reported measures tend to over-report sleep time [39, 40] and under-report awakenings during sleep [40]. Wrist actigraphy has been validated against laboratory-based polysomnography in this age group and shown to have higher accuracy than self-report for sleep duration[41] and awakenings [42]. DXA is a highly accurate method of classifying body tissues and assessing regional body fat distribution [43–45].

This study has some limitations. The sample size was relatively small (n=251). However, it represents 18.6% of the 15-year-old population of Reykjavik in 2015 (n=1355) [46]. The cross-sectional nature precludes study of the temporal relationships between sleep, physical activity, and metabolic factors. All measurements were collected from spring until early summer, a period of drastic change in day length and weather in Iceland which could affect sleep timing [47] and physical activity level [47, 48]. Earlier analyses of this cohort found no association between sleep duration and day length [28] but a positive association between physical activity and day length [29]. We attempted to mitigate the influence of day length by statistically controlling for it in all regression models. Finally, our sample is mostly lean and racially and ethnically homogeneous, potentially limiting the generalizability of the results to other populations.

## Conclusion

Greater nightly variation in sleep duration, lower sleep quality, and less physical activity was associated with a less favorable metabolic profile in 15-year-old adolescents, highlighting the importance of physical activity and maintaining a regular sleep schedule. Further research is needed to determine the longitudinal relationship between sleep, physical activity, and metabolic health from adolescence into adulthood.

## Acknowledgments

The authors would like to thank the participants of the study, the staff at participating schools and the MS students involved in the data collection. They also thank the Icelandic Heart Association, The Icelandic Centre for Research (RANNIS) and the University of Iceland Research Fund for financial support.

